# A Bayesian Perspective on Accumulation in the Magnitude System

**DOI:** 10.1101/101568

**Authors:** Benoît Martin, Martin Wiener, Virginie van Wassenhove

**Affiliations:** CEA, DRF/I2BM, NeuroSpin; INSERM, U992, Cognitive Neuroimaging Unit; Université Paris-Sud; Université Paris-Saclay, F-Gif/Yvette, France; Department of Psychology, George Mason University, Fairfax, VA USA

**Keywords:** space, time, number, decision-making

## Abstract

Several theoretical and empirical work posit the existence of a common magnitude system in the brain. Such a proposal implies that manipulating stimuli in one magnitude dimension (e.g. duration in time) should interfere with the subjective estimation of another magnitude dimension (e.g. size in space). Here, we asked whether a generalized Bayesian magnitude estimation system would sample sensory evidence using a common, amodal prior. Two psychophysical experiments separately tested participants on their perception of duration, surface, and numerosity when the non-target magnitude dimensions and the rate of sensory evidence accumulation were manipulated. First, we found that duration estimation was resilient to changes in surface and numerosity, whereas lengthening (shortening) the duration yielded under- (over-) estimations of surface and numerosity. Second, the perception of surface and numerosity were affected by changes in the rate of sensory evidence accumulation, whereas duration was not. Our results suggest that a generalized magnitude system based on Bayesian computations would minimally necessitate multiple priors.

## INTRODUCTION

The representation of space, time, and number is foundational to the computational brain^1–3^, yet whether magnitudes share a common (conceptual or symbolic) format in the brain is unclear. Walsh’s A Theory Of Magnitude (ATOM)^3^ proposes that analog quantities are mapped in a generalized magnitude system which entails that space, time, and number may share a common neural code. One additional implication for the hypothesis of a common representational system for magnitudes is that the estimation of magnitude in a target dimension (e.g., size in space) should be affected by the manipulation of the magnitude in another non-target dimension (e.g., duration in time), such that the larger the magnitude of the non-target, the larger one should perceive the target magnitude dimension to be (Figure 1A). Such predictions can be formalized in Bayesian terms^4^ so that the magnitude of each dimension yields a likelihood estimate subsequently informed by an amodal prior common to all magnitude dimensions (Figure 1B). In line with ATOM and the common magnitude system hypothesis, a growing body of behavioral evidence^5–29^, for review see^30–32^ suggests the existence of interferences across magnitude dimensions. Several neuroimaging studies also suggest the possibility of a common neural code for quantity estimations mostly implicating parietal cortices^33–39^ but see^28,40^. However, while a variety of interactions between time, space and number has been reported, the directionality of these interactions is not always consistent in the literature^13,14^ suggesting the need to moderate the claim for a common magnitude system: for instance, manipulating the duration of events has seldom been reported to affect numerical and spatial magnitudes^12,13,26,29^ whereas numerosity^6^ and size^5,7^ typically influence duration. Yet, using a literal interpretation of ATOM^3^, if time, number and space shared a common representational system and amodal prior, all magnitude dimensions should interact with each other in a bi-directional manner (Figure 1A).

**Figure 1:**
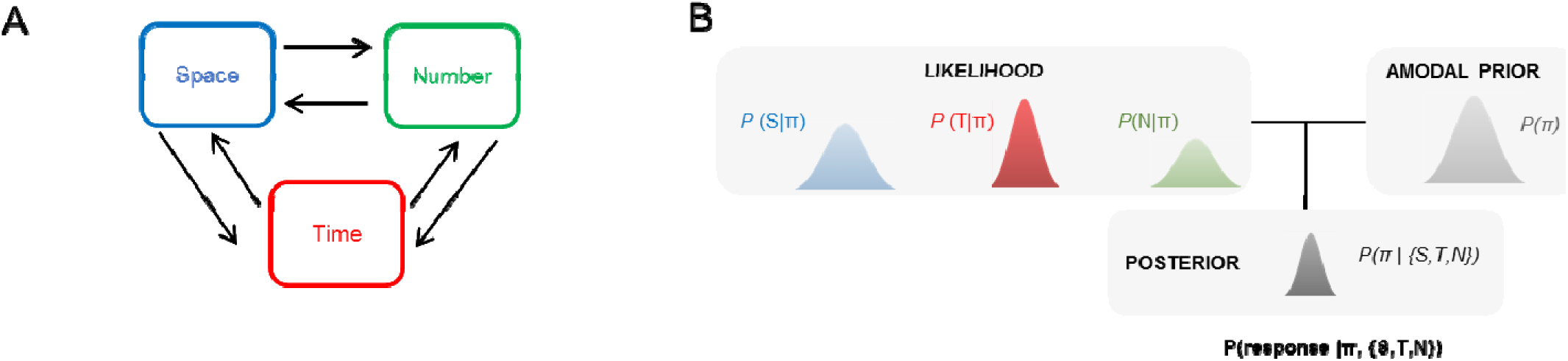
A Bayesian Magnitude System. **A:** Proposals for a common magnitude system in the brain suggest that estimating one magnitude dimension (e.g. space) should be affected by the manipulation of any other magnitude dimensions (e.g. time) (Walsh, 2003). Such interactions should show bidirectional interferences so that, in our example, manipulating the spatial dimension of an event should affect the estimation of its duration in a comparable manner as manipulating its time dimension would affect the estimation of its spatial dimension. **B:** To account for bidirectional interferences across magnitudes in a Bayesian framework for magnitude estimation, a common global or amodal prior can be posited to constrain the estimations of all magnitude dimensions [4]. Under such model, increasing (decreasing) the value of one magnitude should increase (decrease) the estimation of another magnitude. This panel is only for illustration purposes and do not reflect the actual distributions of the magnitudes used in the study.

Recent discussions in the field suggest that the combination and evaluation of quantities in a common representational system would be realized on the basis of Bayesian computations^13,41^. Convergent with this proposal, recent examinations of Bayesian processing in magnitude estimation have demonstrated a number of distinct effects^4^. One primary example is the so-called central tendency effect, wherein magnitude estimates regress to the mean of the stimulus set, such that large (small) magnitudes are under (over) estimated. Crucially, central tendency effects have been demonstrated across a number of different magnitude judgments, including duration (historically known as Vierordt’s law^41,42^, numerosity^27^, distance and angle^43^. Further, correlations between the degree of central tendency have been found between the magnitude of different dimensions^43^, suggesting the existence of “global” priors for common magnitude estimation^4^. The notion of global priors is compatible with a literal read of Walsh’s ATOM model by suggesting that, though differences in the initial processing of different magnitude dimensions may exist, the representation of these magnitudes is amodally stored. Further, the existence of global priors would provide an explanation for congruency effects between different magnitude dimensions. However, a single global prior for magnitude estimations would not explain why congruency effects may be inconsistent between dimensions or why the directionality of interferences may differ across magnitude dimensions.

To address this first working hypothesis, we used a paradigm in which stimuli consisted of clouds of dynamic dots characterized by the total *duration* of the trial (D), the total *number* of dots (N) and the overall *surface* filled by the dots (S) cumulating over the trial. The duration, the number and the cumulative surface of the dots will be thereafter used in reference to the magnitude of the time, number and space dimensions. Two experiments were conducted (Figure 2A-B). In a first experiment (Experiment 1), while participants estimated the magnitude of a target dimension (e.g. D), we independently manipulated the magnitude of non-target dimensions (e.g. S and N). This design allowed us to test all possible combinations and investigate possible interactions between magnitudes (Table 1). If the magnitude of different dimensions interact, increasing or decreasing the N or the S should lead to an under/overestimation of D (Figure 2D). These effects should be bidirectional so that when participants estimate N or S, increasing or decreasing the magnitude of non-target dimensions D should lead to under- or overestimation of the target magnitude dimension N or S. On the other hand, if dimensions are independent, manipulating the number of events in a given trial should not affect duration or surface estimates. Similarly, decreasing or increasing the duration should not affect numerical or spatial judgments if magnitudes are independent.

**Table 1.**
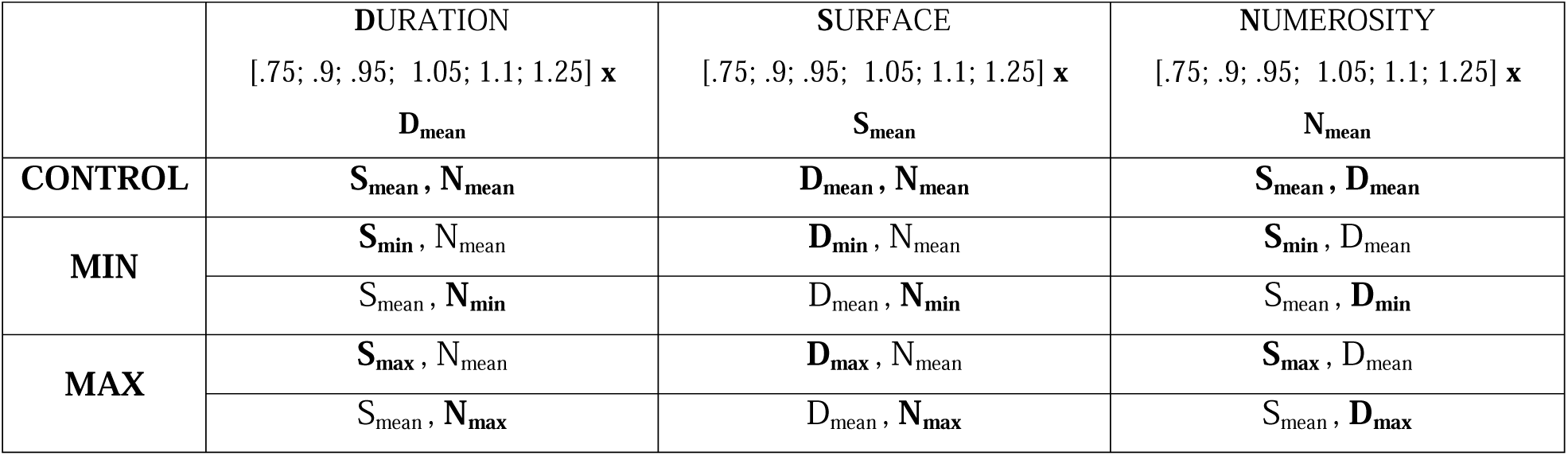
Full design of Experiment 1 testing the interactions across magnitudes with a linear sensory evidence accumulation regime. The tested magnitude dimension could take 6 possible values corresponding to 75, 90, 95, 105, 110 and 125 % of its mean value. In CONTROL trials (first row) of the duration condition (D, second column), participants estimated the duration of the trial when D varied between the 6 possible values while the surface and number dimensions were kept to their mean values (S_mean_ and N_mean_, respectively). In MIN trials (second row), participants estimated the magnitude of a given dimension (e.g. D) varying between the 6 possible values while one of the two non-target magnitude dimensions was kept at its mean value (e.g. N_mean_) and the other was set to its minimal value (e.g. S_min_). In MAX trials (third row), participants estimated the magnitude of a given dimension varying between the 6 possible values while one of the two non-target dimensions was kept at its mean value and the other was set to its maximal value. In total, 72 trials per experimental condition were collected (i.e. 12 trials per tested magnitude value in all possible combinations). D: duration; S: surface; N: number; min = minimal; max = maximal.

| | DURATION<br>[.75; .9; .95; 1.05; 1.1; 1.25] x<br>$D_{\text{mean}}$ | SURFACE<br>[.75; .9; .95; 1.05; 1.1; 1.25] x<br>$S_{\text{mean}}$ | NUMEROSITY<br>[.75; .9; .95; 1.05; 1.1; 1.25] x<br>$N_{\text{mean}}$ |
| --- | --- | --- | --- |
| CONTROL | $S_{\text{mean}}, N_{\text{mean}}$ | $D_{\text{mean}}, N_{\text{mean}}$ | $S_{\text{mean}}, D_{\text{mean}}$ |
| MIN | $S_{\text{min}}, N_{\text{mean}}$ | $D_{\text{min}}, N_{\text{mean}}$ | $S_{\text{min}}, D_{\text{mean}}$ |
| | $S_{\text{mean}}, N_{\text{min}}$ | $D_{\text{mean}}, N_{\text{min}}$ | $S_{\text{mean}}, D_{\text{min}}$ |
| MAX | $S_{\text{max}}, N_{\text{mean}}$ | $D_{\text{max}}, N_{\text{mean}}$ | $S_{\text{max}}, D_{\text{mean}}$ |
| | $S_{\text{mean}}, N_{\text{max}}$ | $D_{\text{mean}}, N_{\text{max}}$ | $S_{\text{mean}}, D_{\text{max}}$ |

**Figure 2:**
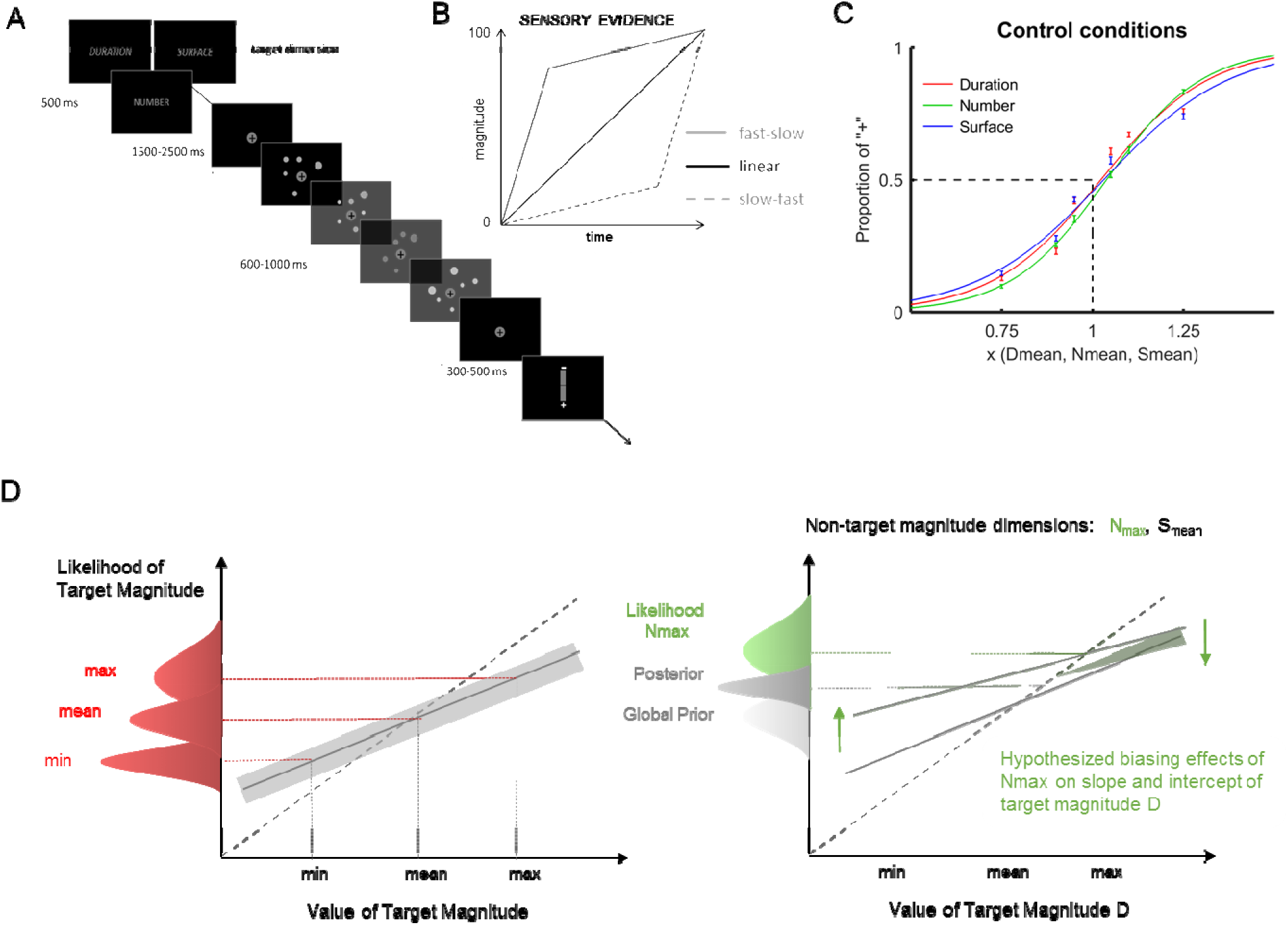
Experimental Design. **A:** On a given trial, participants were presented with a word (“durée”, “nombre” or “surface”) indicating the dimension to estimate. In Experiment 1, one magnitude dimension could vary +/- 25 %, 10 %, and 5% of its mean value, while the second one was set to its minimal or maximal value, and the third one to its mean value (Table 1). At the end of the stimulus presentation, participants used a vertical scale to estimate the target magnitude. **B**: Three distributions were used for evidence accumulation: while D linearly accumulates over time (black trace), the rate of dot presentation could be manipulated to control N and S. Experiment 1 tested a linear distribution (filled black trace); Experiment 2 tested two distributions: a fast-slow (filled grey trace) and a slow-fast (dotted grey trace) distribution. The different stimulus distributions can be experienced with the videos Linear, FastSlow and SlowFast in Supp. Material. **C**: Equated task difficulty across magnitudes. For illustration purposes, the psychometric curve captures the grand average performance obtained for the estimation of Duration, Number and Surface when all non-target dimensions were kept at their mean value. The task difficulty was equated across magnitude dimensions so that no differences in discriminability (PSE and WR) existed between the tested dimensions. Bars are 2 s.e.m. **D:** Predictions for the effect of non-target manipulations on the estimation of the target magnitude dimension. Left panel: varying the target magnitude while keeping the non-target dimensions to their mean values provided the control central tendency and intercept. In a common Bayesian magnitude estimation system [4], comparable tendency and intercept should be predicted pending controlled matching between magnitudes and task difficulty (panel C). Right panel: estimation of D while N is set to its maximal value (in green, Nmax). Maximal value in non-target magnitude may affect the central tendency and the intercept if an amodal global prior common to D and N is implicated in the estimation of duration. In this example, N_max_ would bias the lowest (highest) duration values towards larger (smaller) values and lead the intercept to move upwards.

In a second working hypothesis, we manipulated the accumulation regime of sensory evidence for the estimation of N and S (Figure 2B). The accumulation of sensory evidence in time for space and number has seldom been controlled for or manipulated during magnitude estimations. In a prior experiment^13^, constraining the duration of sensory evidence accumulation in the S and N dimensions, the estimation of duration remained resilient to changes in the other dimensions, whereas D affected the estimation of S and N: curiously, the longer (shorter) durations decreased (increased) the estimation of S and N. These results were discussed in the context of a possible Bayesian integration of magnitudes across dimensions. Similarly, here, using a dynamic paradigm in which N and S accumulate over time raises the question of the implications of varying the speed or rate of sensory evidence delivery: for a given N or S, if D increases, the speed of presentation decreases, and vice versa. Hence, while the number of dots and the cumulative surface accumulated linearly in time in Experiment 1 (Figure 1B), in Experiment 2, we investigated whether changes in the rate of presentation of visual information affected the estimation of D, N, and S. Two evidence accumulation regimes were tested: a fast-slow (FastSlow) and a slow-fast distribution (SlowFast), see Stimuli part in Material & Methods section.

In a third question, we wished to investigate the extent to which Bayesian models could explain the behavioral results obtained in magnitude estimation, independent of the means by which participants provided their estimates. Thus far, studies demonstrating central tendency effects^42–44^ have all relied on continuous estimation procedures, wherein participants estimated a particular magnitude value with a motor response. In particular, these studies utilized reproduction tasks, which required participants to demarcate where (when) a particular magnitude matched a previously presented standard. In contrast, the majority of studies demonstrating congruency effects in magnitude estimation have all employed two-alternative forced choice (2AFC) designs. This difference may be particularly relevant as recent studies have demonstrated that the size-time congruency effect, one of the most heavily studied and replicated, depends on the type of decision being made^45^ (but see^25,46^ for congruency effects with temporal reproduction). As such, for both Experiment 1 and 2, we provide systematic quantifications of the magnitude estimates as categorical estimations together with analysis of continuous reports.

## MATERIALS & METHODS

### Participants

A total of 45 participants were tested. 3 participants did not come to the second session and 10 were disregarded for poor performance: 10 participants either never reach 50% of “+” responses for the largest values, or the p-values associated with the goodness-of-fits in the control conditions were >.05 (see Procedure). Hence, a total of 17 healthy volunteers (7 males, 10 females, mean age 24.9 ± 5.8 y.o.) took part in Experiment 1, and 15 participants (8 males, 7 females, mean age 26.5 ± 7 y.o.) in Experiment 2. All had normal or corrected-to-normal vision. Both experiments took place in two sessions one week apart. Prior to the experiment, participants gave a written informed consent. The study was conducted in agreement with the Declaration of Helsinki (2008) and was approved by the Ethics Committee on Human Research at Neurospin (Gif-sur-Yvette, France). Participants were compensated for their participation.

### Stimuli

The experiment was coded using Matlab 8.4 with Psychtoolbox (v 3.0.12) and built on a published experimental design (see^13^). Visual stimuli were clouds of grey dots which appeared dynamically on a black computer screen (1024 x 768 pixels, 85 Hz refresh rate). Dots appeared within a virtual disk of diameter 12.3 to 15.2 degrees of visual angle; no dots could appear around the central fixation protected by an invisible inner disk of 3.3 degrees. Two dots could not overlap. The duration of each dot varied between 35 ms and 294 ms, and their diameter between 0.35 and 1.14 degrees. A cloud of dots was characterized by its duration (D: total duration of the trial during which dots were presented), its numerosity (N: cumulative number of dots presented on the screen in the entire trial) and its surface (S: cumulative surface covered by the dots during the entire trial). On any given trial, D, N and S could each take 6 possible values corresponding to 75, 90, 95, 105, 110 and 125 % of the mean value. We fixed D to 800 ms (D_mean_ = 800 ms) and initially picked N_mean_ = 30 dots and S_mean_ = 432 mm^2^. The initial values of N_mean_ and S_mean_ were then individually calibrated in the calibration session of the experiment (see Procedure). To ensure that luminance could not be used as a cue to perform the task, the relative luminance of dots varied randomly across all durations among 57, 64, 73, 85, 102 and 128 in the RGB-code.

In Experiment 1, the total number of dots accumulated linearly over time (see Linear video), 2 to 7 dots at a time in steps of 9 to 13 iterations (Figure 2A). In Experiment 2, the total number of dots accumulated in a fast-to-slow or in a slow-to-fast progression: in FastSlow, 75% ± 10% of the total number of dots in the trial were presented in the first 25% of the duration of the trial, whereas in SlowFast, 25% ± 10% of the total number of dots was shown during 75% of the total duration of the trial (Figure 2B; see FastSlow and SlowFast videos).

### Procedure

Participants were seated in a quiet room ~60 cm away from the computer screen with their head maintained on a chinrest. The main task consisted in estimating the magnitude of the trial along one of its three possible dimensions (D, N, or S). Each experiment consisted of two sessions: in the first or calibration session, stimuli were calibrated to elicit an identical discrimination threshold in all three dimensions on a per individual basis (see^13^) and the main objective of the first session was to calibrate an individual’s N_mean_ and S_mean_ with the chosen D_mean_ in order to match task difficulty across dimensions. D_mean_ was kept constant for all participants. The second experimental session consisted in the experiment proper.

In the calibration session of Experiment 1 and 2, the task difficulty across magnitudes was individually calibrated by computing the participant’s Point of Subjective Equality (PSE: 50% discrimination threshold) and the Weber Ratio (WR) for each dimension D, N, and S. The PSE traditionally provides an estimate of an individual’s perceptual threshold: here, the PSE specifically corresponded to the magnitude value in the target dimension at which participants responded at chance level. The WR provided the estimate of the steepness of the fitted psychometric curve, and thus of an individual’s perceptual sensitivity in discriminating magnitudes of the target dimension. A smaller (larger) WR indicates a steeper (flatter) curve and a better (worst) sensitivity. Participants were passively presented with exemplars of the minimum and maximum value for each dimension and were then required to classify 10 of these extremes as minimum ‘-’ or maximum ‘+’ by pressing ‘h’ or ‘j’ on an AZERTY keyboard, respectively. Participants then received feedback indicating the actual number of good answers they provided. Subsequently, the PSE and the WR were independently assessed for each magnitude dimension by varying the magnitude in one dimension and keeping the magnitude in the other two dimensions at their mean values (e.g. if D varied among its 6 possible values, S was S_mean_ and N was N_mean_). 5 trials per magnitude value of the target dimension were collected yielding a total of 30 trials (5 trials x 6 values) per dimension from which the individual’s PSE and WR could be computed and compared. This process (~15 min) was iterated until the individual’s PSE for the target dimension was stable and the WR similar across dimensions. The PSE were considered as matching across dimensions when all of them were between 95 and 105% of the mean magnitude value. For each participant, the mean of the three WR was also calculated, and the WR were considered as matching when: (1) the three WR were in the mean (SD = 2), (2) when the ratio between the larger and the smaller WR was < 3 and (3) each WR value was smaller than 0.25. The final mean magnitude values for each dimension were N_mean_ = 32 (SD = 3 dots) and S_mean_ = 476 (SD = 58 mm^2^) for Experiment 1 and N_mean_ = 32 (SD = 2 dots) and S_mean_ = 490 (SD = 41 mm^2^) in Experiment 2.

In the experimental session of both Experiment 1 and 2, participants were first tested again with 30 trials calibrated to their individual N_mean_ and S_mean_ calculated in the calibration session to ensure that their PSE and WR remained identical so that task difficulty was balanced across magnitude dimension. Only two participants in Experiment 1 required the recalibration procedure to be performed again. Subsequently, participants performed the magnitude estimation task *proper* in which participants were asked to provide a continuous magnitude estimation of the target dimension by moving a cursor on a vertical axis whose extremes were the minimal and maximal magnitude values of the target dimension. In a given trial, participants were provided with the written word ‘Durée’ (Duration), ‘Nombre’ (Number) or ‘Surface’ (Surface) which indicated which target magnitude dimension they had to estimate (Figure 2A). At the end of a trial, the vertical axis appeared on the screen with the relative position of ‘+’ and ‘-’ pseudo-randomly assigned to the extreme bottom or the top of the axis. The cursor was always initially set in the middle position on the axis. Participants used the mouse to vertically move the slider along the axis and made a click to validate their response. They were asked to emphasize accuracy over speed. Trials were pseudo-randomized across dimensions and conditions.

In Experiment 1, five experimental conditions were tested per dimension: in the control condition, the two non-target dimensions were kept to their mean magnitude values; in the four remaining conditions, one of the other non-target dimension was set to its minimal or maximal value, while the other was kept to its mean value (Table 1). A total of 1080 trials were tested in Experiment 1 (3 dimensions x 5 conditions x 6 magnitude values x 12 repetitions).

In Experiment 2, two main sensory accumulation regimes were tested (FastSlow, and SlowFast) and the emphasis was on the effect of D on S and N. The main control condition consisted in assessing the estimation of duration with S_mean_ and N_mean_, and in testing whether the rate of sensory evidence delivery affected the estimation of duration. Two control conditions investigated the effect of the rate of stimulus presentation on the estimation of S and N without varying non-target dimensions. In light of the results obtained in Experiment 1, Experiment 2 did not investigate the interactions of N or S on D, nor the interactions between N and S. Ten experimental blocks alternated between FastSlow and SlowFast presentations counterbalanced across participants. 12 repetitions of each possible combination were tested yielding a total of 144 trials for D (2 distributions x 6 durations x 12 repetitions), 432 trials for N (2 distributions x 3 conditions x 6 numerosities x 12 repetitions) and 432 trials for S (2 distributions x 3 conditions x 6 surfaces x 12 repetitions) for a grand total of 1008 trials in Experiment 2 (Table 2).

**Table 2.**
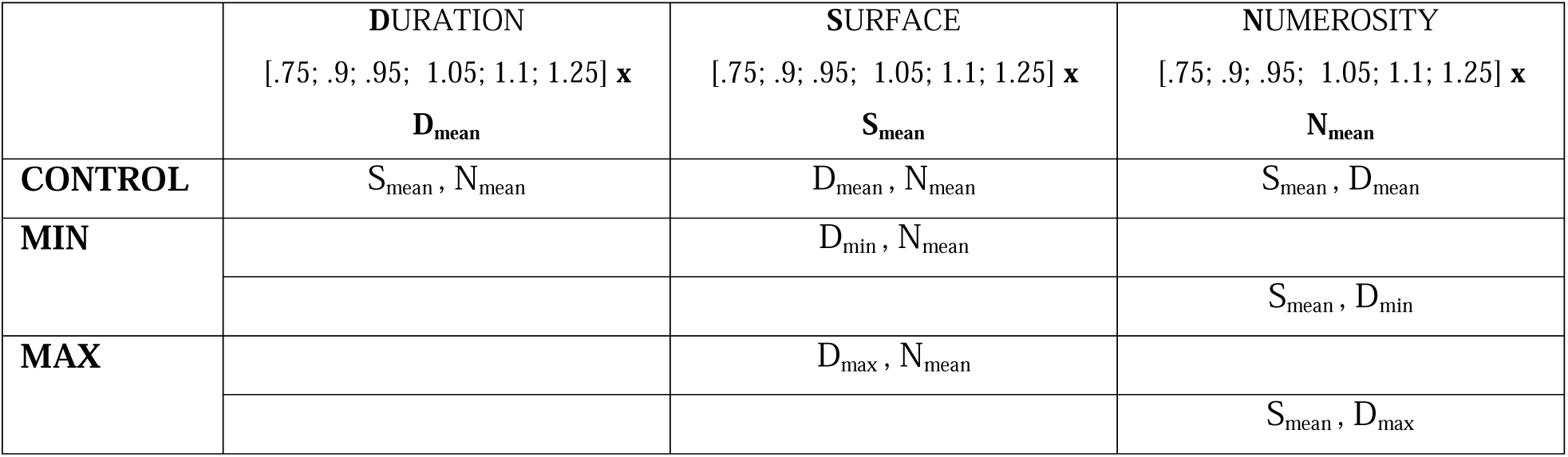
Full design of Experiment 2 testing the effects of duration and FastSlow and SlowFast sensory evidence accumulation regime on magnitude estimation. In CONTROL trials, as in Experiment 1, participants estimated the value of each target magnitude dimension when non-target dimensions were kept to their mean values. In MIN and MAX trials, participants estimated S or N when D was the shortest (D_min_) or the longest (D_max_) and the other dimension was kept to its mean value (N_mean_ and S_mean_, respectively). Importantly in Experiment 2, the clouds of dots could accumulate over time according to two accumulation regimes (FS:FastSlow, and SF:SlowFast). A total of 144 trials per experimental condition was collected (i.e. 12 trials for a given magnitude value in a specific condition x 2 distributions).

|  | DURATION<br>[.75; .9; .95; 1.05; 1.1; 1.25] x<br>D<sub>mean</sub> | SURFACE<br>[.75; .9; .95; 1.05; 1.1; 1.25] x<br>S<sub>mean</sub> | NUMEROSITY<br>[.75; .9; .95; 1.05; 1.1; 1.25] x<br>N<sub>mean</sub> |
| --- | --- | --- | --- |
| CONTROL | S<sub>mean</sub> , N<sub>mean</sub> | D<sub>mean</sub> , N<sub>mean</sub> | S<sub>mean</sub> , D<sub>mean</sub> |
| MIN |  | D<sub>min</sub> , N<sub>mean</sub> |  |
|  |  |  | S<sub>mean</sub> , D<sub>min</sub> |
| MAX |  | D<sub>max</sub> , N<sub>mean</sub> |  |
|  |  |  | S<sub>mean</sub> , D<sub>max</sub> |

### Statistical Analyses

To analyze the point-of-subjective equality (PSE) and the Weber Ratio (WR), participants’ continuous estimates were first transformed into categorical values: a click between the center of the axis and the extreme demarcating ‘+’ (‘-’) was considered as a ‘+’ (‘-’) response. Proportions of ‘+’ were computed on a per individual basis and separately for each target dimension and each experimental condition. Proportions of ‘+’ responses were fitted using the logit function (Matlab 8.4) on a per individual basis. Goodness-of-fits were individually assessed and participants for whom the associated p-values in the control conditions were >.05 were excluded from the analysis. On a per condition basis, PSE and WR that were 2 standard deviations away from the mean were disregarded and replaced by the mean of the group.

This procedure affected a maximum of 2 values per condition across all individuals. Statistics were run using R (Version 3.2.2). PSE and WR were defined as:

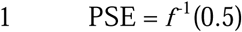

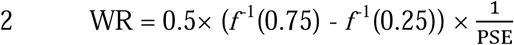

where *f* is the logit function used to fit individuals’ responses, and PSE is the magnitude value of the target dimension when the proportion of ‘+’ responses is equal to 0.5. Using the inverse of the *f* function, the WR was calculated as the mean difference between the just-noticeable-differences (aka magnitude values at 25% and 75% performance) normalized by the individual’s PSE. Additionally, to specifically address central tendency effects, continuous estimates were analyzed. For each magnitude dimension, continuous estimates were expressed as the relative position on the slider that participants selected on each given trial, with higher percentages indicating closer proximity to ‘+’. To measure the central tendency effects, continuous estimates were plotted against the corresponding magnitude dimension for each condition, also expressed as a percentage – where 0 indicated the smallest magnitude and 100 indicated the largest – and fits with a linear regression, and the slope and y-intercept of the best fitting line were extracted^47,48^. Slope values closer to 1 indicated veridical responding (participants responded with perfect accuracy), whereas values closer to 0 indicated a complete regression to the mean (participants provided the same estimate for every magnitude). In contrast, intercept values of these regressions could indicate an overall relative bias for over- or under-estimation so that higher (lower) intercept values would indicate that participants overestimated (underestimated) the magnitude in the target dimension^49^. To compare central tendency effects between magnitude dimensions, correlation matrices between slope values for D, N, and S were constructed. Bonferroni corrections were applied to control for multiple comparisons.

## RESULTS

To examine Bayesian effects in the magnitude system, we evaluated both categorical and continuous judgments in two magnitude estimation experiments using variations of the same paradigm (Figure 2A). To evaluate choice responses, continuous estimates were binned according to which end of the scale they were closer to. Previous work has demonstrated that bisection tasks and continuous estimations are compatible and provide similar estimates of duration^44,50^. Our intention was thus to first replicate the effects of Lambrechts and colleagues^13^ with a modified design, and second, to measure central tendency effects in our sample to examine whether these effects correlated between magnitude dimensions, which would suggest the existence of global priors (see^4^, Figure 2D).

### Control conditions: matching task difficulty across magnitude dimensions

Two independent repeated-measures ANOVAs with the PSE or WR as dependent variables using magnitude dimensions (3: D, N, S), control conditions (3: Linear (Experiment 1), SlowFast and FastSlow (Experiment 2) distributions as within-subject factors did not reveal any significant differences (all *p* >.05). This suggested that participants’ ability to discriminate the different values presented in the tested magnitudes was well matched across magnitude dimensions (Figure 2C).

#### Experiment 1: Duration affects the estimation of Number and Surface

We first analyzed the data of Experiment 1 as categorical choices. Figure 3A illustrates the grand average estimations of duration, numerosity, and surface for all experimental manipulations (colored traces) along with changes of PSE (insets).

**Figure 3:**
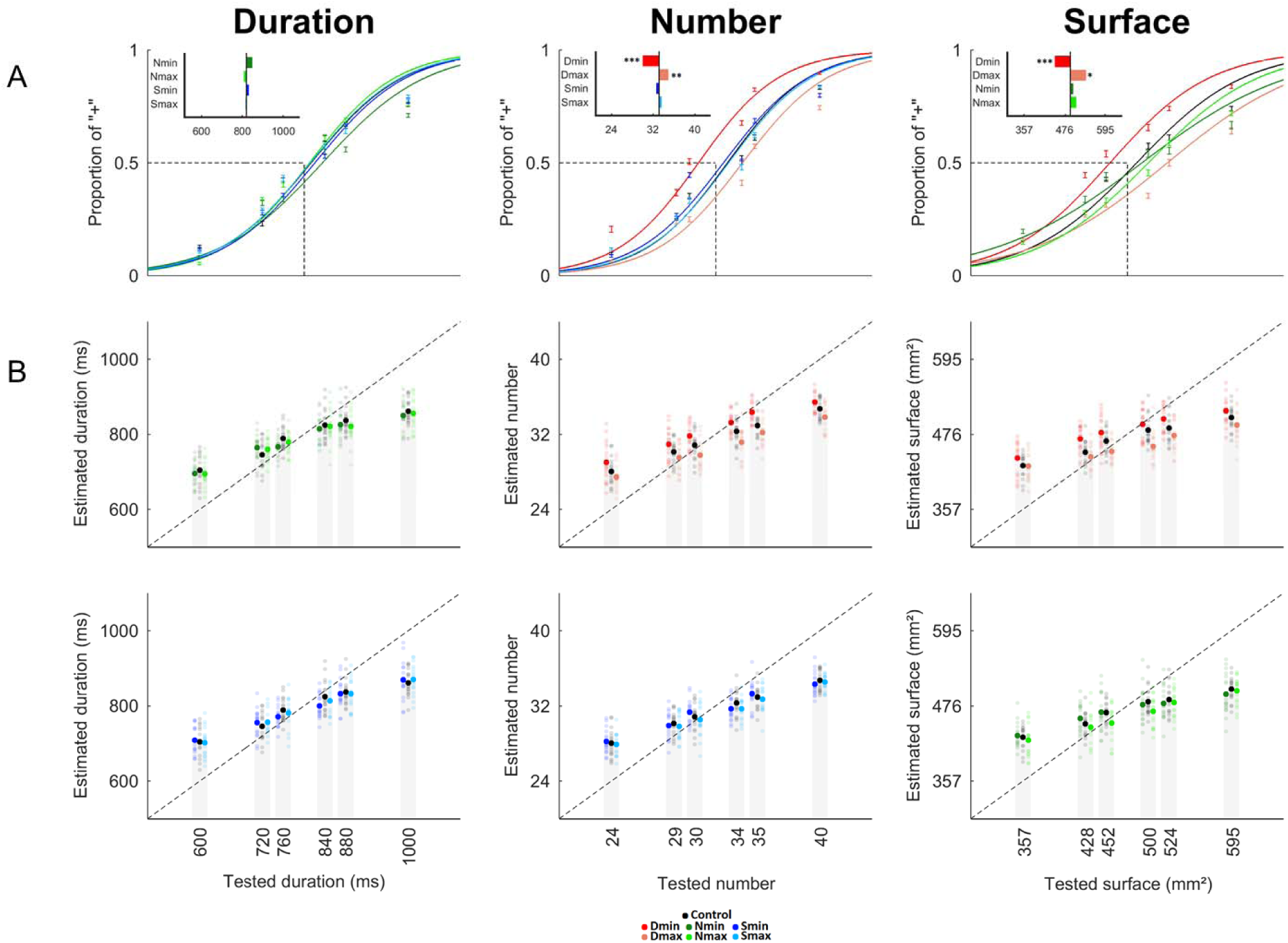
Duration affects the estimation of S and N (Experiment 1). **A**: Categorical quantifications and PSE. The percentage of « + » responses as a function of the target magnitude dimension (D: left panel, N: middle panel, S: right panel) were fitted on a per individual basis. The inset in the top left of each figure depicts the shift in PSE for each experimental condition compared to the control condition represented by the black vertical line. No effects of N or S on D were found; no effects of S on N or of N on S were found; both N and S were significantly overestimated when presented during the shortest duration (Dmin, red) and underestimated when presented with the longest duration (Dmax, orange). **B**: Continuous judgments. Individual performances (transparent dots) and mean performances (filled dots) for the estimation of D (left panels) with changing N (green) and S (blue); of N (middle panels) with changing D (red) and S (blue); and of S (left panels) with changing D (red) and N (green). The continuous scale was mapped from 0 to 100. The dotted line is the ideal observer’s performance. All experimental conditions showed a central tendency. No effects (intercept or central tendency) of N or S on D were found (left top and bottom graphs, respectively); no effects of S on N (middle bottom graph) or of N on S (left bottom graph) were found. Significant main effects of D were found on the central tendency and the intercept of N (middle top) and S (left top). N_min_: miminal numerosity value; N_max_: maximal numerosity value; S_mm_: minimal surface value; S_max_: maximal surface value; D_min_: minimal duration value; D_max_: maximal duration value. *** p< 0.001; ** p< 0.01; * p< 0.05; bars are 2 s.e.m.

Separate 2x2 repeated-measures ANOVAs were run on the PSE obtained during the magnitude evaluation of each target dimension using the non-target dimensions (2) and their magnitude values (2: min, max) as within-subject factors. No main effects of non-target dimension (F[1,16] = 0.078, *p* = 0.780), magnitude value of the non-target dimension (F[1,16] =0.025, *p* =0.875) or their interaction (F[1,16] = 0.003, *p* =0.957) were found on duration (D) indicating that manipulating N or S did not change participants’ estimation of duration (Fig. 3A, left panel). In the estimation of N, main effects of non-target dimensions (F[1,16] = 7.931, *p* = 0.0124), their magnitudes (F[1,16]=25.53, *p* =0.000118) and their interactions (F[1,16] = 23.38, *p* =0.000183) were found. Specifically, when the magnitude of the non-target dimensions were at their minimal value, the PSE obtained in the estimation of N was lower than when the magnitude of the non-target dimensions were at their maximal value. Additionally, in the estimation of N, D_min_ lowered the PSE more than S_min_, and D_max_ raised the PSE more than S_max_. Paired t-tests were run contrasting the PSE obtained in the estimation of N during the control (D_mean_ S_mean_) and other experimental conditions: D_min_ significantly increased [PSE(D_min_) < PSE(D_mean_): *p* = 4.1e^-5^] whereas D_max_ significantly decreased [PSE(D_max_) > PSE(D_mean_): *p* = 0.0032] the perceived number of dots (Fig. 3A, middle panel, inset). There were no significant effects of S on the estimation of N. Altogether, these results suggest that the main effect of changing the magnitude in the non-target dimension on numerosity estimation was driven by the duration of the stimuli.

In the estimation of S, we found no main effect of the non-target dimension (F[1,16] = 1.571, *p* = 0.228) but a significant main effect of the magnitude values in non-target dimensions (F[1,16]=22.63, *p* =0.000215). The interaction was on the edge of significance (F[1,16] = 3.773, *p* =0.0699) suggesting that, as for N, only the magnitude of one non-target dimension may be the main driver of the significant results observed in the effect. Paired t-tests contrasting the PSE obtained in the estimation of S during the control (Dmean N_mean_) and other conditions showed that D_min_ significantly increased (PSE(D_min_) < PSE(D_mean_): *p* = 8.7e^-4^), whereas D_max_ significantly decreased (PSE(D_max_) > PSE(D_mean_): *p* = 0.035) the perceived surface (Fig. 3A, right panel). No significant effects of N on S were found. As observed for the estimation of N, these results suggest that the main effect of non-target magnitude on the estimation of S was entirely driven by the time dimension.

Overall, the analysis of PSE indicated that participants significantly overestimated N and S when dots were presented over the shortest duration, and underestimated N and S when dots accumulated over the longest duration. Additionally, manipulating N or S did not significantly alter the estimation of duration. No significant interactions between N and S were found. To ensure that these results could not be accounted for by changes in participants’ perceptual discriminability in the course of the experiment, repeated-measures ANOVA were conducted independently for each target dimension (3: D, N, S) with the WR as dependent variable and the experimental conditions (5) as main within-subject factors. No significant differences (all *p* > .05) were found suggesting that the WRs were stable over time, and that task difficulty remained well matched across dimensions in the course of the experiment.

For the analysis of continuous estimates (Figure 3B), we first examined the effect of central tendency for each target dimension, collapsing the data across the non-target dimensions (Figure 4A). A repeated measures ANOVA of slope values with magnitude as a within-subjects factor revealed a main effect of the magnitude of the target dimension [F(2,32) = 13.284, *p* = 0.000063]. Post-hoc paired t-tests identified this effect as driven by a lower slope, indicating a greater regression to the mean for S as compared to D [*t*(16) = 3.495, *p* = 0.003] and N [*t*(16) = 5.773, *p* = 0.000029], with no differences in slope values between D and N [*t*(16) = 0.133, *p* = 0.896] (Figure 4B).

**Figure 4:**
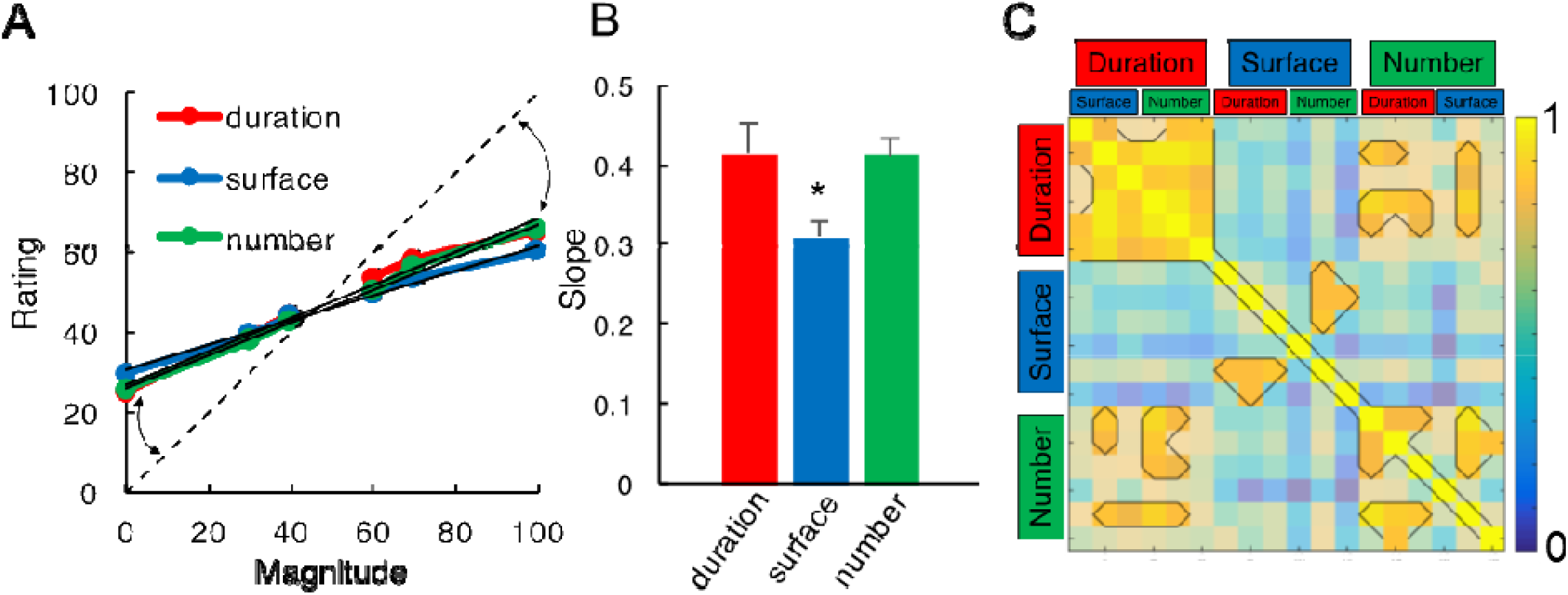
Central Tendency Effects in Magnitude Estimation (Experiment 1). A: Average continuous estimates for all three magnitude dimensions as target, collapsed across all non-target magnitude dimensions. Magnitudes were normalized as a percentage of the maximum presented magnitude value, with zero representing the smallest and 100 representing the largest magnitude presented. Continuous estimates were similarly normalized as a percentage of the sliding scale, with zero representing a minimal estimate of “-” and 100 representing a maximal estimate of “+”. The dashed identity line indicates where estimates should lie for veridical performance. Deviations from this identity line (arrows) exhibited central tendency, wherein smaller and larger magnitudes were over- and under-estimated, respectively. **B:** Slope values extracted from a best fitting linear regression in A quantify the degree of central tendency, with smaller values indicating greater regression to the mean. Similar slope values were observed for duration and number, but significantly lower estimates were found for surface estimates than either duration or number. **C:** Correlation matrix of slope values for every target and non-target magnitude trial type. Each target magnitude dimension was tested in the presence of three possible values (min, mean, max) of a non-target dimension. Outlined pixels represent those Pearson correlation coefficients that survived a multiple comparison correction. Slope values across non-target dimensions were correlated within each target dimension (sections along the diagonal); crucially, slope values for duration and number were correlated with each other when each one was the target magnitude (lower left and upper right sections). Further, no correlation between surface and either duration or number as target dimensions were observed (middle top and bottom sections). * indicates p<.05; bars are 2 s.e.m.

Further analyses revealed comparable findings as in the categorical analysis: separate 2x3 repeated measures ANOVAs were run for each magnitude dimension, with the non-target dimension and its magnitude value as within-subject factors. Analysis of slope values revealed no significant main effects or interactions for any of the tested magnitudes (all *p* >.05), indicating no change in central tendency, as a function of the non-target magnitudes. However, an analysis of intercept values demonstrated a significant main effect of the non-target dimension for S [*F*(2,32) = 24.571*,p* < 0.00001] and N [*F*(2,32) = 39.901*,p* < 0.00001], but not for D [*F*(2,32) = 0.010*,p* = 0.99]. Specifically, intercept values were shifted higher (lower) when D_min_ (D_max_)was the non-target dimension in both the S and N tasks (Figure 3B, red hues), but not when the non-target dimension was N for S, S for N, or for either S or N when D was the target magnitude dimension.

To examine the central tendency effects across magnitude dimensions, we correlated the slope values between target magnitude dimensions (Figure 4C). Collapsing across the non-target dimensions, we found that all three slope values significantly correlated with one another [D to S: Pearson *r* = 0.594; D to N: *r* = 0.896; S to N: *r =* 0.662]. Given that S exhibited a greater central tendency than D or N, we compared the Pearson correlation coefficients with Fisher’s z-test for the differences of correlations; this analysis revealed that the D to N correlation was significantly higher than the D to S correlation [Z = 2.03, *p* = 0.04], and marginally higher than the S to N correlation [Z = 1.73, *p* = 0.083], suggesting that D and N dimensions, which had similar slope values, were also more strongly correlated with each other than with S. To further explore this possibility, we conducted partial Pearson correlations of slope values; here, the only correlation to remain significant was D to N, when controlling for S [*r* = 0.8352], whereas D to S, controlling for N, and N to S, when controlling for D were no longer significant [*r* = 0.0018 and 0.3627, respectively].

The results of the correlation analysis revealed that D and N tasks were highly correlated in slope, indicating that individual subjects exhibited a similar degree of central tendency for these two magnitude dimensions (Figure 4A). From a Bayesian perspective, these results suggest that the priors for D and N may be more correlated than the priors for D and S, or for N and S; thus D, N and S do not share one single prior, but may rather rely on different priors which would be more or less correlated between each other. To explore this at a more granular level, we expanded our correlation analysis to include all non-target dimensions (Figure 4C). The result of this analysis, with a conservative Bonferroni correction (*r* > 0.8) for multiple comparisons confirmed the above results, demonstrating that D and N dimensions were correlated across most non-target dimensions, but that D and N dimensions were weakly and not significantly correlated with S. This finding suggests that D and N estimation may rely on a shared prior, that is separate from S; however, a shared (D, N) prior would not explain why D estimates were unaffected by changes in N, nor would it explain why S estimates are affected by changes in D.

Lastly, we sought to compare the quantifications based on continuous data with those from the categorical analysis. Previous work has demonstrated that the WR on a temporal bisection task correlates with the central tendency effect from temporal reproduction^44^. To confirm this, we measured the correlations between the slope values of continuous magnitude estimates with the WR from the categorical analysis. As predicted, we found a significant negative correlation between slope and WR for D (*r* = −0.69) and N (*r* = −0.57); however, the correlation for S failed to reach significance (*r* = −0.41, *p* = 0.1), indicating that greater central tendency (lower slope values) were associated with increased variability (larger WR). This finding is notable, as the analysis of WR values did not reveal any difference between magnitude dimensions. This suggests that the slope of continuous estimate judgments may be a better measure of perceptual uncertainty than the coefficient of variation derived from categorical responses.

#### Experiment 2: Duration is robust to accumulation rate, not N and S

In Experiment 2, participants estimated D, N or S while the accumulation regime was manipulated as either FastSlow or SlowFast (Fig. 2B, Table 2). As previously, we systematically analyzed the categorical and the continuous reports. First, we tested the effect of the accumulation regime on the estimation of each magnitude dimension by using a 2x3 repeated-measures ANOVA with PSE measured in control conditions (Table 2, 1^st^ row) as independent variable and distribution (2: FastSlow, SlowFast) and magnitude dimension (3: N, D, S) as within-subject factors. Marginal main effects of accumulation regime (F[1,28] = 2.872, *p* = 0.0734) and magnitude dimensions (F[1,14] = 4.574, *p* = 0.0506) were observed. Their interaction was significant (F[2,28] = 10.54, *p* = 0.0004). A post-hoc t-test revealed no significant effects of accumulation regime on the estimation of D (*p* = 0.23), but significant effects of accumulation regime in the estimation of N (*p* =0.016) and S (*p* = 0.0045) (Figure 5A).

**Figure 5:**
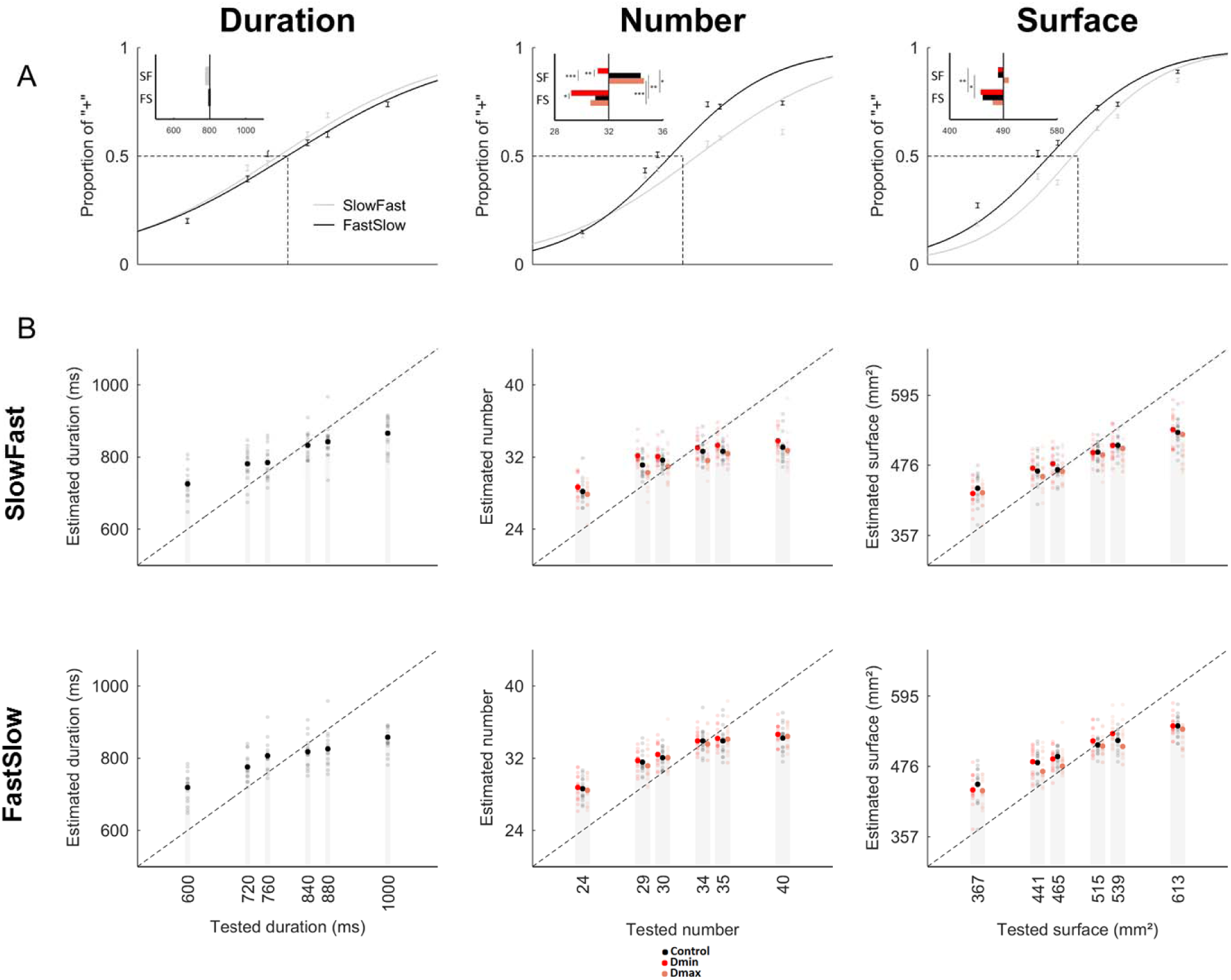
The Accumulation Regime affects Numerosity and Surface but not Duration (Experiment 2). **A:** Psychometric curves illustrate the grand average proportion of “+” as a function of duration (left panel), number (middle panel) or surface (right panel) when the sensory evidence accumulation was manipulated. The SlowFast (SF) results are reported in light gray, the FastSlow (FS) results are reported in black. Left panel: top inset reports the mean PSE of duration estimation observed for each distribution as compared to the ideal observer (vertical black line). Middle panel: top inset reports the mean PSE of number estimation observed for each distribution (SF, FS) and manipulation of D (D_mean_: black, D_min_: red, D_max_: pink). Right panel: top inset reports the mean PSE of surface estimation observed for each distribution (SF, FS) and manipulation of D (D_mean_: black, D_min_: red, D_max_: pink). The estimation of D was not affected by the accumulation regime of sensory evidence whereas N and S were overestimated in the FastSlow as compared to the SlowFast distribution. N and S were also affectd by manipulating of the non-target duration. **B:** Individual (transparent dots) and mean (filled dots) continuous judgments for the estimation of D (left panel), N (middle panel) and S (right panel) with SlowFast (top row) and FastSlow (bottom row) regimes of sensory evidence accumulation. The dotted line is the ideal observer’s performance. D_min_: minimal duration value; D_max_: maximal duration value. *** p< 0.001; ** p< 0.01; * p< 0.05; bars are 2 s.e.m.

Second, we tested the effect of D and accumulation regime on the estimation of N and S (Figure 5A, top insets). We conducted a 2x2x2 repeated-measures ANOVA with PSE as an independent variable and magnitude dimension (2: N, S), accumulation regime (2: FastSlow, SlowFast), and duration (2: D_min_, D_max_) as within-subject factors. Main effects of accumulation regime (F[1,14] = 22.12, *p* = 0.000339) and duration (F[1,14] = 27.65, *p* = 0.000121) were found, suggesting that both N and S were affected by the distribution of sensory evidence over time, and by the duration of the sensory evidence accumulation. No other main effects or interactions were significant although two interactions trended towards significance, namely the two-way interaction between accumulation regime and duration (F[1,14] = 3.482, *p* = 0.0831) and the three-way interaction between dimension, accumulation regime, and duration (F[1,14] = 3.66, *p* = 0.0764). These trends were likely driven by the SlowFast condition as can be seen in Figure 5A.

For the analysis of continuous data (Figure 5B), we first examined any overall differences in slope values for different accumulation regimes (FastSlow *vs.* SlowFast) across all three target magnitude dimensions (D, N, S). A (3x2) repeated measures ANOVA with the above as within-subjects factors revealed a main effect of magnitude dimension [*F*(2,32) = 7.878, *p* = 0.002], with S once again demonstrating the largest slope value, but no effect of accumulation regime or interaction (both *p* >.05), suggesting that the rate of accumulation did not influence the central tendency effect. However, on the basis of our *a priori* hypothesis, post-hoc tests revealed a significantly lower slope value for N in SlowFast compared to FastSlow [*t*(17) = 3.067, *p* = 0.007], suggesting that participants exhibited more central tendency for numerosity when the accumulation rate was slow in the first half of the trial (Figure 5B). The analysis of intercept values did not reveal any effects of accumulation regime or magnitude dimension (all *p* >.05). However, on the basis of our *a priori* hypothesis, post-hoc tests demonstrated that S exhibited a significantly lower intercept for SlowFast compared to FastSlow [*t*(17) = 3.609, *p* = 0.002], with no changes for either D or N (both *p* >.05), indicating that participants underestimated surface when the rate of evidence accumulated slowly in the first half of the trial.

For S and N, further examination of slope values for the three possible durations using a 2x2x3 repeated measures ANOVA with magnitude dimension (2: S, N), accumulation regime (2: FastSlow,SlowFast), and duration (3: D_min_, D_mean_, D_max_) as within-subjects factors, revealed a significant main effect of magnitude dimension [*F*(2,32) = 7.717, *p* = 0.013] and of accumulation regime [*F*(2,32) = 11.345, *p* = 0.004], but not of duration [*F*(2,32) = 1.403, *p* = 0.261]. Using the same analysis for intercept values, we found no main effects of magnitude dimension [*F*(2,32) = 1.296, *p* = 0.272], but a significant effect of accumulation regime [*F*(2,32) = 5.540, *p* = 0.032] and of duration [*F*(2,32) = 21.103, *p* = 0.000001]. More specifically, we found that intercept values were lower for longer durations, indicating greater underestimation when the interval tested was longer. No other effects reached significance (all *p* >.05).

Overall, these findings indicate that duration estimations were immune to changes in the rate of accumulation of non-target magnitudes, similar to the findings of Experiment 1. Also similar, we found that estimates of S and N were affected by duration as non-target magnitude dimension, with longer durations associated with greater underestimation of S and N (Figure 6). In addition, our results demonstrate a difference between accumulation regimes for S and N, with SlowFast regimes associated with greater underestimation than FastSlow, regardless of duration. Lastly, we observed that SlowFast accumulation regimes led to an increase in the central tendency effect, suggesting that slower rates of accumulation may increase reliance on the magnitude priors.

**Figure 6:**
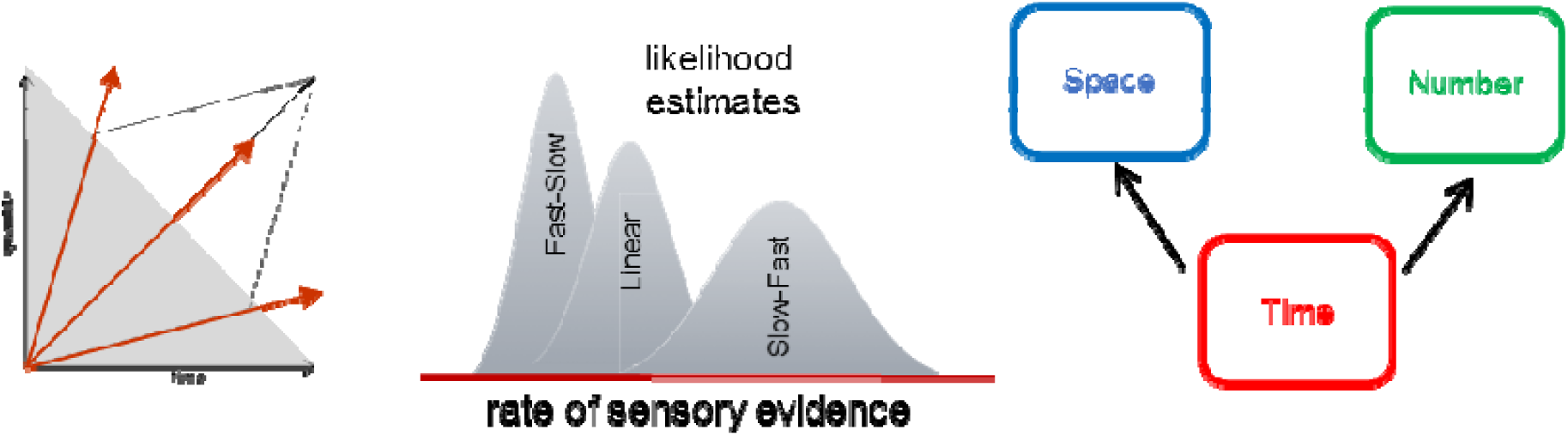
Accumulation regime influences Bayesian estimates. Our results suggest that the speed or rate of sensory evidence accumulation early in the trial (shaded region) and the duration of the trial affect the estimation of surface and numerosity. Additionally, slower rates of sensory evidence are associated with greater uncertainty; greater uncertainty results in increased reliance on the priors. To accommodate these findings, we suggest that the rate of sensory evidence is effectively estimated independently of the total duration although the duration may regulate the noise level in sensory evidence accumulation of quantities.

## Discussion

In this study, we report that when sensory evidence steadily accumulates over time, and when task difficulty is equated across magnitude dimensions (space, time, number), duration estimates are resilient to manipulations of number and surface, whereas number and surface estimates are biased by the temporal properties of sensory evidence accumulation. Specifically, we replicated the findings of Lambrechts and colleagues^13^ by demonstrating that number and surface estimates are under- and overestimated when presented for long and short durations, respectively. These results complement the findings that duration can be resilient to numerosity interference^27^, and that the direction of the interference between space and time may go in opposite direction when using dynamic displays^13^,^28^. Although prior findings have reported asymmetrical effects in magnitude estimation, our findings differ in several ways. First, participants provided a quantified estimation of a given magnitude dimension, allowing a direct assessment of performance within a Bayesian framework in mind (specifically characterizing properties of the central tendency effects as a function of magnitude interferences). In previous reports of interference of number on duration^5,6^, behavioral effects were concluded on the basis of increased reaction times and error rates during incongruent condition presentation (e.g., small number presented with a long duration) which prevented the direct evaluation of participants’ magnitude perception *per se.* As such, no clear direction of interference effects could be concluded from the studies beyond the existence of an interference. Our experimental design also prevented participants to explicitly enumerate the dots in a given trial unlike in previous experiments, in which the speed of dot presentation enabled counting which yielded an influence of N on D^6^.

In a study^7^ using dynamic display which addressed a question close to the ones we address with this experimental approach, participants judged the spatial length or the duration of a growing line. As previously discussed elsewhere (see^13^), the spatial task could have been performed using the coordinates of the line on the screen irrespective of the duration it took the line to grow, possibly explaining why duration was irrelevant for the spatial estimate. In other words, the duration in this task was likely the noisiest cue which in turn was affected by the least noisy cue (i.e. the spatial dimension). Consistent with our current results, a recent study^28^ showed that a longer duration yielded an underestimation of length. In this experiment, the environment was dark (fMRI study) and participants had no access to visual cues to constrain their spatial estimate of the moving dot. In this context, the results showed that the shorter duration increased the distance of the moving dot, consistent with the present findings. Hence, and consistent with previous literature, the lack of robust bidirectional interactions between magnitude dimensions does not support a literal interpretation of ATOM; however, we do not argue that time, number and space do not interact under certain constraints, and rather consider our results to favor a more liberal interpretation of ATOM. Specifically, by considering a Bayesian model relying on multiple priors (one for each dimension), magnitudes may interact in the context of conflicting sensory cues. Recent hypotheses suggest that a Bayesian framework can provide a general explanation for the variety of behavioral features observed in magnitude estimations independently applied to distance, loudness, numerical or temporal judgments^4^. The proposed Bayesian framework combines an estimate of the likelihood (sensory input) with a prior representation (memory). One major goal of our study was thus to determine the degree to which different magnitude dimensions might rely on an amodal global prior representation of magnitude as would be expected using a literal interpretation of a generalized magnitude system such as ATOM^3^. To accomplish this, participants took part in two experiments independently manipulating the congruence across magnitude dimensions (Experiment 1) and the rate of sensory evidence provided to participants (Experiment 2).

A first prediction was that if different magnitude dimensions rely on a single amodal prior, then magnitude estimates should exhibit similar levels of central tendency across magnitude dimensions (duration, surface, number; Figure 2D). Instead, in Experiment 1, our results demonstrated that surface estimates exhibited greater central tendency than either duration or number, and surface estimates were not correlated with the degree of central tendency for either dimension. However, duration and number did exhibit correlated central tendency effects. This finding suggests that estimates of surface are distinct from estimates of duration and number, but that duration and number may be more similar to one another. Indeed, neural recording studies in the prefrontal and parietal cortex of non-human primates have revealed overlapping, yet largely separate, representations of duration and size^51,52^, and number and size^53,54^. Further, while number, size, and time exhibit common activations of the right parietal cortex, they each engage larger networks of regions beyond this area^28,55,56^. For size estimates, recent work suggests that comparisons of object size draw on expectations from prior experience in other brain regions^57^. Yet, as no strong bidirectional effects were observed between duration and number, it is unlikely that duration and number share neuronal populations with similar tuning features.

Another possible interpretation of the results obtained in Experiment 1 is to consider multiple priors in magnitude estimations. When participants make temporal judgements, the combination of prior knowledge P(π) and noisy sensory inputs P(D|π) (duration of the given trial) enables participants to make an accurate posterior estimate, represented by P(π|D) ∞ P(D|π)·P(π) (see^4^, Box 3 for more details), which explains the regression to the mean. Neither numerosity nor surface priors are present in this equation, which could explain why duration estimates are robust to numerosity or surface manipulations. Because numerosity and surface accumulate over time, one possible strategy for the participants was to estimate numerosity and surface based on both the speed of presentation of stimuli, and on the duration of the trial. Specifically, a high (low) speed over a long (short) duration of presentation corresponded to a large (small) value of numerosity or surface. In other words, these results seemed to indicate that, using dynamic displays, the speed of events influences the estimation of magnitudes and yield opposite directionality in the interference across magnitudes. If participants used the speed of event presentation and overestimated (underestimated) D, they also overestimated (underestimated) N or S. As the computation of speed also relies on the duration, both speed and duration become important cues in the estimation process but may have distinct impacts. In Experiment 1, the central tendency effect showed that the shortest duration was overestimated, which could explain why participants overestimated N and S for D_min_; conversely, the longest duration was underestimated, which may explain why participants underestimated N and S. Under this hypothesis, the uncertainty related to the temporal dimension may add noise in the decision or the accumulation process, so that the perceived duration of the trial can bias numerosity and surface estimates (Figure 6). When numerosity and surface accumulate over a given duration, if that duration is short (long) it will affect the accuracy of participants’ estimations. This explanation would be compatible with the hypothesized effect of duration as introducing noise on sensory accumulation, and Experiment 2 was specifically designed to tease out the effect of speed changes in the accumulation rate on the estimation of the target magnitude.

Indeed, one noteworthy aspect unique to the time dimension is that the objective rate of presentation is fixed^58^. That is, objective time by conventional measurements proceeds at a single mean rate. In contrast, we can experimentally manipulate the rate at which we present information for number and surface. In Experiment 1, in order to keep the values of surface and number fixed when duration was manipulated, we necessarily had to change the rate of accumulation for these values. For example, between short and long durations with the same value of number, we had to change the rate of accumulation for number so that the same total value was reached at the end of the duration. This may explain the incongruent effects of duration on surface and number; shorter (longer) durations may engender larger (smaller) estimates of surface and number because the rate of accumulation is faster (slower). In this sense, surface and number are not being influenced by the duration magnitude of the time dimension *per se,* but rather the time dimension is interfering with the rate of accumulation, and so the effect of duration may be an epiphenomenon of the experimental design. To test this hypothesis, we modulated the accumulation rate of the presentation of numerosity and surface in Experiment 2.

In Experiment 2, where the rate of accumulation for number and surface experienced a rate-change a little less (more) than halfway through the presentation time from fast-to-slow (slow-to-fast), we replicated and extended our findings of Experiment 1. Specifically, we again found that shorter (longer) durations led to longer (shorter) estimates of surface and number, regardless of the rate-change in accumulation regime. However, we also found a difference in accumulation regimes: when the rate of accumulation was slower in the beginning of the session, estimates of surface and number were smaller than when the rate of accumulation was faster. It is important to remember that the ultimate value of the presented surface and number was the same, regardless of the accumulation regime. As such, participants were biased in their estimates by the rate of evidence accumulation in the first-half of the given trial, regardless of how long that trial lasted. This strongly suggests that human observers are biased by the rate of accumulation at the start of a trial, and are resistant to changes in rate throughout the trial.

This observation is important in the context of ongoing discussions on drift-diffusion processes in which the accumulation of evidence following the first end-point depends on terminated processes, guess probability (see^59^ case study 1 and Figure 1; see^60^ Figure 2, 4A) and on the importance of change points during the accumulation process^61,62^. Changes in the accumulation rate performed in Experiment 2 imposed a nonlinearity in the accumulation process: the observation that an earlier rate change has a larger impact on magnitude estimation than a later rate change is reminiscent of the ‘primacy effect’ reported in evidence accumulation models, possibly indicative of suppression of newer information by old information^63^. Additionally, this finding strengthens the hypothesis that the effect of duration on surface and numerosity may occur as a result of the impact on the implicit timing or accumulation rate, and not as a function of the explicitly perceived duration. This would be consistent with recent findings suggesting that noise memory - known to scale with duration - was not the primary factor of errors in decision-making but that noise in sensory evidence was instead a major contributor^64^. Our results suggest that speeding up the rate of evidence and lengthening the duration of a trial may be equivalent to increasing noise in sensory accumulation of other magnitude dimensions (Figure 6).

In Experiment 2, we also investigated the effect of accumulation regime on central tendency as participants again provided continuous magnitude estimates on a vertical sliding scale. Previous magnitude studies using continuous estimates have demonstrated a central tendency effect, with over(under)-estimations for small (large) magnitudes^4^, the degree to which depends on the uncertainty inherent in judging the magnitude in question^44,65^. The result of this analysis revealed that, when the rate of accumulation was slow (fast) in the beginning of the trial, the degree of central tendency was greater (lesser); further, the objective duration of the trial did *not* impact central tendency. This finding suggests that slower accumulation regimes engender greater uncertainty in magnitude estimates, and that this uncertainty may be present before the ultimate decisional value is reached. Previous work in decision-making with evidence accumulation has suggested that the objective duration of a trial leads to greater reliance on prior estimates, as longer presentation times are associated with greater uncertainty^66^. Specifically, Hanks and colleagues^66^ found that a drift-diffusion model that incorporated a bias signal to rely on prior evidence that grows throughout the trial could explain reaction time differences in a dots-motion discrimination task. Notably, the bias signal is incorporated into the drift-diffusion process, such that longer trials push the accumulation rate towards a particular value, depending on the prior. A critical manipulation in this study was the emphasis on speed or accuracy for subjects; increased emphasis for accuracy led to longer decision times and greater reliance on the prior, as explained by the model. Our results suggest otherwise – the duration of the trial alone cannot determine reliance on the prior. If the effect of duration solely led to greater reliance on the prior, then we should have seen central tendency effects increase with longer durations, which did not occur in either Experiments 1 or 2. Instead, the rate of evidence accumulation determined reliance on the prior(s), regardless of duration, with slower rates leading to greater reliance.

Additional studies using neuroimaging techniques such as M/EEG need to investigate the neural correlates underpinning accumulation processes in the brain when estimating magnitudes (Centro-Parietal Positivity, for example, see^67,68^), to fully explain the behavioral results obtained in these two experiments. Further, fMRI studies must be conducted to elucidate the neural circuits for memory representations of different magnitudes^48^. Bayesian approaches may provide interesting perspectives on magnitudes estimations, and additional studies need to be performed to understand to which extent these models can be applied to explain the variety of results observed in the literature. One intriguing observation is the finding that duration estimates were not only resilient to changes in numerosity or surface, but also to the rate of sensory evidence. This finding is unexpected and runs counter-intuitive to various findings in time perception. In this task, these robust findings suggest that unlike surface and numerical estimates, duration may not rely on the accumulation of discretized sensory evidence.

## DATA ACCESSIBILITY

Data will be made available through Open Science Framework

## AUTHORS’ CONTRIBUTIONS

B.M., M.W. and V.v.W. designed the study. B.M. collected the data. B.M., M.W. and V.v.W. analyzed the data. All authors co-wrote the manuscript and gave final approval for publication.

## ACKNOWLEDGMENTS

We thank members of UNIACT at NeuroSpin for their help in recruiting and scheduling participants, and Baptiste Gauthier and Tadeusz W. Kononowicz for discussions.

